# Landscape of Dysregulated Placental RNA Editing Associated with Preeclampsia

**DOI:** 10.1101/797365

**Authors:** Xiaoxue Yang, Jing Yang, Xiaozhen Liang, Qian Chen, Sijia Jiang, Haihua Liu, Yue Gao, Zhonglu Ren, Yanhong Yu, Mei Zhong, Xinping Yang

## Abstract

Dysregulated RNA editing is well documented in several diseases such as cancer. The extent to which RNA editing might be involved in diseases originated in the placenta remains unknown, because RNA editing has rarely been studied in the placenta. Here, we have systematically profiled RNA editome on the placentae from 9 patients with early-onset severe preeclampsia (EOSPE) and 32 normal subjects, and a widespread RNA editing dysregulation in EOSPE has been identified. The mis-edited gene set is enriched with known preeclampsia-associated genes and differentially expressed genes in EOSPE. The “RNA editing events” at two microRNA binding sites in 3’-UTR of the *LEP* mRNA were generated, which could lead to increased expression level of *LEP* in trophoblast cells. Upregulation of *LEP* were also observed in the placentae of PE patients. These results suggest that widespread placental RNA editing may be involved in placental development and dysregulation of RNA editing in the placenta may contribute to the pathogenesis of preeclampsia.

## Introduction

RNA editing was first discovered in the mitochondrial cytochrome c oxidase (COX) subunit II gene from trypanosomes in 1986 (Benne et al. 1986). It includes RNA sequence modifications, such as insertion or deletion of nucleotides and nucleotide conversion from cytidine (C) to uridine (U) (Chen et al. 1987; Powell et al. 1987; Benne 1994; Simpson et al. 2004), or from adenine (A) to inosine (I) (Bass and Weintraub 1987; Levanon et al. 2004). The C-to-U RNA editing involves site-specific changes in the mRNAs (Blanc and Davidson 2003), as are well documented in apolipoprotein B (*ApoB*) (Anant and Davidson 2001), neurofibromatosis type 1 (*NF1*) and N-Acetyltransferase 1 (*NAT1*) (Yamanaka et al. 1997; Mukhopadhyay et al. 2002). The most common RNA editing in mammals is A-to-I conversion, which is catalyzed by enzymes called adenosine deaminases acting on RNA (ADARs) (Polson et al. 1991; Savva et al. 2012). A-to-I RNA editing widely occurs in Alu transposons and noncoding regions such as introns and untranslated regions (UTRs) and in coding regions as well (Eisenberg and Levanon 2018).

RNA editing can lead to missense codon changes (Nishikura 2016), alternative splicing (Rueter et al. 1999), modification of microRNAs (miRNAs) and long noncoding RNAs (lncRNAs) (Kawahara et al. 2007; Gong et al. 2017), and miRNA binding sites (Nishikura 2010). The functions of RNA editing in noncoding regions are based on the site-specific modifications of RNA motifs, which mediate interactions with other molecules. Accumulating evidence on the function of RNA editing points to the regulation of RNA expression levels (Sommer et al. 1991; Nishikura 2010). RNA editing may regulate RNA expression levels through directly mediating RNA degradation (Kim et al. 2005), indirectly through alternative splicing (Rueter et al. 1999; Laurencikiene et al. 2006; Hsiao et al. 2018), or through modification of miRNA binding sites (Wang et al. 2013; Nakano et al. 2016). The suggested role for RNA-editing-mediated RNA degradation is supported by the observation that the editing enzyme ADAR interacts with UPF1, a mediator of nonsense-mediated decay (NMD) (Agranat et al. 2008). RNA editing sites are enriched in intronic regions in sequencing data of RNA population purified by poly-d(T) (Huntley et al. 2016), suggesting that there might be an association between RNA editing and intron retention. Intron retention can affect gene functions in ways such as bringing in new functional elements (Buckley et al. 2011), down-regulating RNA expression through NMD (Belgrader et al. 1994), and interfering mRNA translation (Middleton et al. 2017). Several studies demonstrate that RNA editing in UTRs regulates mRNA degradation through disrupting miRNA binding (Wang et al. 2013; Nakano et al. 2016; Nakano et al. 2017).

RNA editing has been found in several human diseases (Maas et al. 2006), such as cancer and neurodegeneration (Dominissini et al. 2011; Singh 2012; Han et al. 2015; Hwang et al. 2016), involving in pathways of apoptosis (Wang et al. 2004; Liu et al. 2009), innate immune response (Mannion et al. 2014), cellular proliferation and DNA repair (Yeo et al. 2010). Genome-scale studies on RNA editing sites in cancer have revealed the critical roles of RNA editing in the global regulation of gene expression (Fumagalli et al. 2015; Han et al. 2015; Paz-Yaacov et al. 2015). Trophoblast cells in the placenta have some features similar to cancer cells (Holtan et al. 2009), and the placenta is a vital organ in human development, yet RNA editing in the placenta has been rarely documented. We do not know to what extent RNA editing might be involved in the pregnancy complications with placenta origin, such as preeclampsia (PE). Here, we carried out RNA editome profiling on placentae from 9 patients with early-onset severe preeclampsia (EOSPE) and 32 normal controls, and found 2,920 differentially edited sites (DESs) located in 1,229 genes. The genes with dysregulated RNA editing were enriched with known PE-associated genes, and also with differentially expressed genes (DEGs) in EOSPE. We further generated two mutations to mimic “RNA editing events” at two miRNA binding sites in 3’-UTR of *LEP*, leading to increased expression level of *LEP*. This is consistent with the observed upregulation of *LEP* in the placenta of EOSPE patients. These results suggest that widespread RNA editing dysregulation in placentae might be involved in the pathogenesis of EOSPE.

## Results

### RNA editome profiling

We performed RNA editome profiling on placental transcriptomes of 9 patients with EOSPE and 32 normal controls (**Supplemental Table S1**), using an in-house pipeline to search for PE-associated RNA editing sites (**Supplemental Fig. S1**). After removing adaptors and low quality reads, the cleaned reads were aligned to human reference genome (hg38 version) by tophat2 (**Supplemental Table S2**) and reliable variants were detected using Genome Analysis Toolkit (GATK) (McKenna et al. 2010; Kim et al. 2013). We identified 629,721 and 1,233,453 variants in EOSPE and normal samples respectively. After removing the variants that failed to pass the variant quality score recalibration (VQSR) filtering and/or had allele frequency higher than 0.05 in East Asian population (Eas_AF > 0.05) (See Methods), we detected 142,372 variants in 9 EOSPE samples and 399,565 variants in 32 normal samples respectively. Among these variants, 236,569 were identified as A/G or C/T mismatches, corresponding to A-to-I or C-to-U conversions in RNA (**Fig. 1A and 1B**), which are two main forms of RNA editing in mammals. Most of the potential RNA editing sites were located in transposons, especially in Alu transposons (**Fig. 1C**), consistent with previous studies in cancers (Bazak et al. 2014). About 89% (210,406/236,569) of these variants were located in noncoding regions (introns and intergenic regions) (**Fig. 1D**). The A-to-I RNA editing was the main editing form in most of the genomic regions, while C-to-U editing was found more often in exons.

Figure 1. RNA editome of placentae from patients with EOSPE and from normal subjects. (A) Flowchart for identification of RNA editing sites in placentae from EOSPE patients and normal subjects. Red boxes represent the analyses; blue boxes represent results of the analyses. EOSPE: early-onset severe preeclampsia; DES: differentially edited site; UTR: untranslated region. See Figure S1 for detailed flowchart. (B) RNA editing types in placentae. (C) Distribution of RNA editing sites in transposon elements (TE) and non-transposon regions. Transposon elements were extracted from RepeatMasker Database. (D) Distribution of RNA editing sites in exons, introns, 5’-UTRs, 3’-UTRs and intergenic regions. The genomic regions were defined by RefGene annotation. Numbers above the bars are the number of the sites. (E) Data overlaps and differences between our placental RNA editome dataset and RNA editing databases. (F) Validation of 89 RNA editing sites by Sanger sequencing and multiple alignment of the reads. Yellow bars represent the fraction of validated RNA editing sites and blue bars represent the fraction of genomic DNA mutations. Numbers within the bars are the number of the sites (all validation results of 89 sites were listed in **Supplemental Table S3**).

There are rarely studies on RNA editing in the placenta, and even the most comprehensive database REDIportal does not cover placenta (Picardi et al. 2017). Thus, the placenta-specific RNA editing sites are depleted in current RNA editing databases. We compared the RNA editing sites we discovered in the placenta with known RNA editing sites in the DARNED (Kiran and Baranov 2010), RADAR (Ramaswami and Li 2014) and REDIportal databases (Picardi et al. 2017). There are 3,149 RNA editing sites in our dataset overlapping with all of the three RNA editing databases, and about 48% (112,868/236,569) of placental RNA editing sites are recorded in REDIportal database. The remaining 121,749 RNA editing sites are not included in any of these three databases, indicating that these RNA editing sites may be placenta specific (**Fig. 1E**). To empirically validate our dataset, we randomly picked 33 RNA regions with 89 RNA editing sites and amplified the genomic DNA regions by PCR for Sanger sequencing (**Supplemental Table S3**). The validation rates were 80% for exonic editing sites, 71% for intronic editing sites and nearly 100% for editing sites in UTR regions (**Fig. 1F**). These results demonstrated that our placental editome dataset is of high-quality.

### Differentially edited sites (DESs) are associated with EOSPE and gene expression level

For each RNA editing site, we calculated the editing ratio in every sample and evaluated the editing ratio difference between EOSPE and normal placenta by one-tailed Wilcoxon test (See Methods). At the cutoff *P =* 0.05, we obtained 2,920 differentially edited sites (DESs) which exhibited significant difference of editing ratio between EOSPE and normal subjects (**Fig. 2A and Supplemental Table S4**).

The genes with DESs were significant enriched for PE-associated genes collected from literature (**Fig. 2B and Supplemental Table S5**), and for differentially expressed genes (DEGs) in EOSPE (**Fig. 2C, Supplemental Table S6** and related manuscript by Ren et al.), suggesting the DESs may be involved in PE. To explore the biological functions and pathways of the genes with DESs, we performed enrichment analysis of these genes in Gene Ontology (GO) terms and Kyoto Encyclopedia of Genes and Genomes (KEGG) pathways using ClusterProfiler (Yu et al. 2012). We found that the genes with DESs are significantly enriched in terms such as the female pregnancy, placenta development and autophagy (adjusted *P <* 0.05, **Supplemental Fig. S2A and Supplemental Table S7**), and that the genes with DESs are significantly enriched in nine pathways, including endocytosis, focal adhesion, ubiquitin mediated proteolysis and protein processing in endoplasmic reticulum (**Fig. 2D and Supplemental Table S7**).

Figure 2. Overview of EOPSE differentially edited sites (DESs). (A) Distribution of DESs in genomic regions. Genomic regions were defined by RefGene annotation. Numbers above the bars are the numbers of the sites. See Table S4 for all DESs. (B-C) Fraction of known PE-associated genes (B) and fraction of differentially expressed genes (DEGs) of EOSPE (C) in the genes with DESs. *P*-values were calculated by Fisher-exact test in R. *: *P <* 0.05; **: *P <* 0.01; ***: *P <* 0.001; ****: *P <* 0.0001; n.s.: no significance. The standard error of the fraction was estimated using a bootstrapping method with 10,000 resamplings. PE-associated genes and EOSPE DEGs were listed in Supplemental Table S5 and S6. (D) The KEGG pathways enriched with genes harboring DESs. Color of bars: adjusted *P*-values; X-axis: the number of genes harboring DESs in the pathways. Enrichment analysis and *P*-value calculation were carried out using ClusterProfiler. (E) Gene-pathway-bipartite network for genes with DESs. Yellow diamonds represent the KEGG pathways, and red circles represent up-regulated genes in EOSPE, blue down-regulated genes and gray genes with no significant expression change. See Supplemental Table S7 for *P-*values for pathway enrichment (D) and gene-pathway interactions (E).

The endocytosis pathway contains 30 genes with DESs, including 5 EOSPE up-regulated genes (*ARPC2, PARD6B, EPS15, RHOA* and *EGFR*) and 3 EOSPE down-regulated genes (*ARAP3, RAB11FIP1* and *CYTH3*) (**Fig. 2E**), suggesting that the endocytosis pathway may be regulated by differential editing and may be involved in the pathogenesis of EOSPE. The focal adhesion pathway contains 23 genes with DESs, including 4 EOSPE up-regulated genes (*FLT1, DIAPH1, RHOA* and *EGFR*) (**Fig. 2E**). The gene *FLT1* is one of the few known potential biomarkers of PE (Thomas et al. 2007; Jebbink et al. 2011; McGinnis et al. 2017), and the genes *RHOA* and *EGFR* are also reported to be associated with PE (Hornsby et al. 1989; Faxen et al. 1998; Zhou and Qiao 2006), suggesting that the focal adhesion function may be commonly affected in the placenta of EOSPE.

To investigate if these DESs were associated with gene expression levels, we classified all placenta samples into two groups, with vs. without RNA edited sites (edited group and not-edited group), for each DES (See Methods). One-tailed Wilcoxon test was applied to examine the difference in the expression levels of the gene containing the editing site between edited and not-edited group. We found significant differential expression for 352 genes with 542 DESs (*P* < 0.05, **Table S8**). We also calculated the correlation coefficient between editing ratios and the corresponding gene expression levels for DESs, and found that 305 DESs showed significantly correlation between editing ratio and gene expression level (**Supplemental Table S8**), suggesting that the editing ratio for some editing sites on the transcripts of a gene might regulate the gene expression level.

### DESs in exons, introns and 3’-UTRs are associated with PE

Of the 2,920 DESs, 225 were located in exons, 1,621 in introns, and 335 in 3’-UTRs. We calculated the enrichment of the literature-curated PE-associated genes (**Supplemental Table S5**) and DEGs in EOSPE (**Supplemental Table S6**) in the corresponding genes of these DESs. The genes harboring DESs in the three different regions were enriched with PE-associated genes respectively, suggesting the disease-association of DESs in each category (**Fig. 3A**). The genes with intronic DESs were also enriched with DEGs in EOSPE (**Fig. 3B**), suggesting that intronic DESs might be involved in the dysregulation of gene expression in this disease. Figure 3. DESs in exons, introns and 3’-UTRs. (A-B) Fraction of known PE-associated genes (A) or DEGs of EOSPE (B) in genes with DESs in different regions. *P*-values were calculated by Fisher-exact test in R. *: *P <* 0.05; **: *P <* 0.01; ***: *P <* 0.001; ****: *P <* 0.0001; n.s.: no significance. The standard error of the fraction was estimated using a bootstrapping method with 10,000 resamplings. (C) Mutation types of exonic DESs. Mutation types were annotated based on RefGene using ANNOVAR. The numbers above the bars are the numbers of DESs. (D) Association between intronic DESs and gene expression. Red circle of the Venn diagram represents intronic DESs with significant gene expression difference, and yellow circle represents intronic DESs with significant correlation between editing ratio and gene expression level. See Supplemental Table S8 for all the intronic DESs associated with gene expression. (E) Bipartite network of miRNAs and genes with DESs in 3’-UTR. The miRNA-targeting regions harboring DESs in 3’-UTRs were predicted by TargetScan. Yellow: genes with DESs in 3’-UTRs; blue: miRNAs.

Among the 225 exonic DESs, 108 were located in protein-coding genes and 117 in noncoding RNAs. Of the 108 protein-coding exonic DESs, 73 (18 A-to-I and 55 C-to-U) were synonymous, 31 (16 A-to-I and 15 C-to-U) were nonsynonymous, 3 were stop-gain and 1 was unknown (**Fig. 3C**). Among these 34 nonsynonymous/stopgain DESs, 26 exhibited up-regulated editing levels in EOSPE samples (**Supplemental Fig. S3A**). Eight of the nonsynonymous DESs were predicted to have damaging effects on proteins according to the SIFT and Polyphen2 values by ANNOVAR (**Table 1**). For example, the DES at the position chr3:49360951 in exon of *RHOA* was predicted as a damaging mutation. *RHOA* showed increased editing ratio in EOSPE samples and had interactions with multiple DEGs of EOSPE (**Supplemental Fig. S3A and S3B**), suggesting that this DES in *RHOA* may play a role in the development of EOSPE.

**Table 1.** Exonic differentially edited sites that predicted as damaging/harmful

| Chr<sup>a)</sup> | Pos<sup>b)</sup> | Type | Gene | Avsnp147 | SIFT<br>Pred | Polyphen2<br>Pred |
| --- | --- | --- | --- | --- | --- | --- |
| chr1 | 111476845 | C-to-U | C1orf162 | rs150674974 | D<sup>c)</sup> | B<sup>d)</sup> |
| chr1 | 155612848 | C-to-U | MSTO1 | rs622288 | D | B |
| chr3 | 49360951 | A-to-I | RHOA | - | D | - |
| chr3 | 122341557 | C-to-U | CSTA | rs34173813 | D | P<sup>e)</sup> |
| chr3 | 195780494 | C-to-U | MUC4 | rs75737861 | D | B |
| chr10 | 45826300 | C-to-U | AGAP4 | rs201626545 | D | B |
| chr10 | 46383535 | C-to-U | ANXA8L1 | - | - | P |
| chr10 | 93681515 | C-to-U | FRA10AC1 | rs11187583 | D | P |

<sup>a)</sup>Chr, chromosome; <sup>b)</sup>Pos, position; <sup>c)</sup>D, damaging; <sup>d)</sup>B, benign; <sup>e)</sup>P, possibly damaging

It is reported that RNA editing tends to occur in introns (Athanasiadis et al. 2004), and some intronic RNA editing sites are known to regulate gene expression by influencing circRNA biogenesis and alternative splicing (Rueter et al. 1999; Laurencikiene et al. 2006; Ivanov et al. 2015; Hsiao et al. 2018). We speculated that there might be an association between intronic DESs and intron retention (IR). We identified IR events by comparing the transcripts assembled by StringTie with the canonical transcripts from GENCODE (Pertea et al. 2016) (See Methods). For every IR event, we calculated the normalized percent spliced in (PSI) and compared the PSI between edited group and not-edited group (See Methods). We identified 359 intronic DESs in the retained introns, and found that 36 IR events with 42 intronic DESs which exhibited significant PSI difference between edited and not-edited group by one-tailed Wilcoxon test (**Supplemental Fig. S3C**). Most of these DESs exhibited increased PSI in the edited group. Among these 42 DESs, 17 exhibited significantly difference of gene expression between edited and non-edited group. These results show that some intronic DESs are associated with intron retention.

To evaluate the association of the intronic DESs and the expression levels of corresponding genes, we compared the expression difference of the genes between two groups of placental samples with differential RNA editing (See Methods). We observed 390 intronic DESs located in genes which exhibit significant differential expression between placental samples with vs. without RNA edited sites (edited group vs. not-edited group) (**Fig. 3D and Supplemental Table S8**). We further calculated the correlation coefficient between RNA editing ratio and gene expression level, and found that 211 sites showed significant correlation between editing ratio and the expression level of corresponding genes (**Fig. 3D and Supplemental Table S8**), suggesting that these intronic DESs may regulate the gene expression level.

As a common regulatory strategy used by the cell, microRNA (miRNA) can bind to specific sequences in the 3’-UTRs of target mRNAs and regulate their stability (Wang et al. 2013; Nakano et al. 2016; Nakano et al. 2017). We predicted 2,219 miRNA-targeting regions in genes harboring DESs using TargetScan (Shi et al. 2017), and 51 DESs are located in 45 known miRNA-targeting regions (**Fig. 3e and Supplemental Table S9**). After discarding the DESs located in regions with null preferentially conserved targeting (PCT) values, we identified 31 DESs that may regulate miRNA binding (**Table 2**).

**Table 2.** 31 DESs in 3’-UTRs that located in miRNA-targeting regions with non-null PCT^a)^ values.

| MiRNA | Chr<sup>b)</sup> | Position | Gene | Transcript |
| --- | --- | --- | --- | --- |
| miR-10-5p | chr1 | 35602159 | PSMB2 | ENST00000373237.3 |
| miR-219-5p | chr1 | 52824220 | ZYG11B | ENST00000294353.6 |
| miR-24-3p | chr1 | 100024500 | SLC35A3 | ENST00000465289.1 |
| miR-22-3p | chr1 | 202585129 | PPP1R12B | ENST00000608999.1 |
| miR-150-5p | chr1 | 204552462 | MDM4 | ENST00000391947.2 |
| miR-455-3p.2 | chr1 | 220058113 | BPNT1 | ENST00000322067.7 |
| miR-10-5p | chr2 | 98623012 | MGAT4A | ENST00000264968.3 |
| miR-375 | chr2 | 147931120 | ORC4 | ENST00000392857.5 |
| miR-214-5p | chr7 | 17344771 | AHR | ENST00000242057.4 |
| miR-9-5p | chr7 | 128255810 | LEP | ENST00000308868.4 |
| miR-9-5p | chr7 | 128255811 | LEP | ENST00000308868.4 |
| miR-10-5p | chr8 | 1782334 | CLN8 | ENST00000331222.4 |
| miR-184 | chr8 | 43023708 | HOOK3 | ENST00000307602.4 |
| miR-217 | chr11 | 32853772 | PRRG4 | ENST00000257836.3 |
| miR-217 | chr11 | 78069179 | NDUFC2 | ENST00000281031.4 |
| miR-214-5p | chr12 | 50930723 | METTL7A | ENST00000332160.4 |
| miR-214-5p | chr14 | 21460463 | RAB2B | ENST00000397762.1 |
| miR-214-5p | chr16 | 10529809 | EMP2 | ENST00000359543.3 |
| miR-141-3p/200a-3p | chr16 | 13239082 | SHISA9 | ENST00000558583.1 |
| miR-217 | chr16 | 13239082 | SHISA9 | ENST00000558583.1 |
| miR-490-3p | chr19 | 2836037 | ZNF554 | ENST00000317243.5 |
| miR-196-5p | chr19 | 4654589 | TNFAIP8L1 | ENST00000327473.4 |
| let-7-5p/98-5p | chr19 | 4654589 | TNFAIP8L1 | ENST00000327473.4 |
| miR-214-5p | chr19 | 16550610 | SLC35E1 | ENST00000595753.1 |
| miR-214-5p | chr19 | 51519173 | SIGLEC6 | ENST00000346477.3 |
| miR-17-5p/20-5p/93-5p/106-5p/519-3p | chr19 | 52584380 | ZNF701 | ENST00000540331.1 |
| miR-192-5p/215-5p | chr19 | 52839344 | ZNF468 | ENST00000595646.1 |
| miR-214-5p | chr19 | 57861909 | ZNF587 | ENST00000339656.5 |
| miR-216a-5p | chr20 | 33646143 | CBFA2T2 | ENST00000375279.2 |
| miR-216b-5p | chr20 | 33646143 | CBFA2T2 | ENST00000375279.2 |
| miR-10-5p | chrX | 2906938 | ARSD | ENST00000381154.1 |
| miR-216a-5p | chrX | 24077208 | EIF2S3 | ENST00000253039.4 |
| miR-216b-5p | chrX | 24077208 | EIF2S3 | ENST00000253039.4 |
| miR-26-5p | chrX | 77827544 | MAGT1 | ENST00000358075.6 |
| miR-455-3p.2 | chrX | 77828504 | MAGT1 | ENST00000358075.6 |

<sup>a</sup>PCT: preferentially conserved targeting; <sup>b</sup>Chr: chromosome

### RNA editing in 3’-UTR regulates *LEP* expression by disrupting miRNA binding

Of all the DESs in 3’-UTRs, 24% (81/335) show significant correlation (*P <* 0.05) between editing ratio and gene expression level (**Table S8**), and five of them, including *EIF2S3*:chrX:24077208:A-to-I, *LEP*:chr7:128255810:A-to-I, *LEP*:chr7:128255811:A-to-I, MGAT4A:chr2:98623012:A-to-I and *ZNF554*:chr19:2836037:A-to-I, are located in the predicted miRNA-targeting regions (**Fig. 4A and Table 2**). The *LEP* gene is known to be highly up-regulated in preeclamptic placenta (Enquobahrie et al. 2008), and the elevated circulating leptin level is an indicator for severe preeclampsia (Teppa et al. 2000). In our EOSPE placental transcriptome, the *LEP* expression exceeded normal level by more than 60 folds. We identified 11 DESs, and 10 of them are located in 3’-UTR of *LEP* (**Fig. 4B**). We evaluated the expression level of *LEP* and *miR-9-5p* in the placenta samples from 9 EOSPE patients and 9 randomly picked normal subjects using quantitative real-time PCR (qRT-PCR). The *LEP* was significantly up-regulated in the EOSPE compared to the normal controls (**Fig. 4C**), whereas *miR-9-5p* showed no significant expression difference (**Fig. 4D**). This missing correlation between *LEP* and *miR-9-5p* expression level indicated that the change of *LEP* expression level in patients was not due to a *miR-9-5p* expression change.

Figure 4. DESs in 3’-UTRs regulate gene expression by inhibiting miRNA binding. (A) Association of DESs in 3’-UTRs with gene expression level. Samples were classified into edited group or not-edited group based on the existence of the RNA editing event at a given editing site. The relative expression level was estimated as CPM (counts per million bases) using edgeR. One-tailed Wilcoxon test was used to evaluate the gene expression difference between edited group and not-edited group using R package. *LEP.1* referred to ‘*LEP*:chr7:128255810:A-to-I’; *LEP.2* referred to: ‘*LEP*:chr7:128255811:A-to-I’. See Table S8 for CPM values of these sites. (B) Two clusters of DESs (C1 and C2) in *LEP* 3’-UTR were shown in red letters and labeled as S1-S10. S6 and S7 were located in the miRNA-targeting region and thus were chosen for testing the effect of RNA editing on miRNA binding. (C-D) The qRT-PCR results of *LEP* (C) and *miR-9-5p* (D) on placentae of 8 EOSPE patients and 8 randomly-chosen normal subjects. *ACTB* (C) or *U6* (D) was taken as internal control. Wilcoxon test was performed to evaluate the difference of expression levels between EOSPE and normal placentae using R package. Error bars represent the standard errors. (E) The qRT-PCR results of *LEP* in HTR-8/SVneo cells with/without *miR-9-5p* transfection. *ACTB* was used as internal control. Wilcoxon test was performed to evaluate the difference of *LEP* expression levels using R package. Error bars represent the standard errors. (F) The results of luciferase assay on HTR-8/SVneo cells transfected with plasmids carrying edited *LEP* 3’-UTR (*LEP*_*MUT2-UTR*_) or not-edited *LEP* 3’-UTR (*LEP*_*WT-UTR*_) at S6 and S7 sites. Y-axis represents the relative luciferase activity. Wilcoxon test was used to evaluate the difference of luciferase activity using R package. Error bars represent the standard errors. Significance in (A) and (C-F): *: *P <* 0.05; **: *P <* 0.01; ***: *P <* 0.001; ****: *P <* 0.0001; n.s.: no significance. See Supplemental Table S10 for numerical values in (C-F). (G) Schematic to illustrate the mechanism for 3’-UTR RNA editing in the regulation of *LEP* expression.

Among the 10 DESs in the 3’-UTR of *LEP*, 2 sites (chr7:128255810:A-to-I, S6 and chr7:128255811:A-to-I, S7) (**Fig. 4B**) are located in *miR-9-5p* targeting region, and the editing ratios of these two DESs are significantly correlated with the expression level of *LEP* (**Supplemental Table S8**). These results suggest that these two DESs in *LEP* 3’-UTR may alter *miR-9-5p* targeting motif and thus may inhibit miRNA-induced *LEP* degradation, which may in turn lead to an upregulation of *LEP*.

To test this hypothesis, we first evaluated the regulation of *miR-9-5p* on *LEP* expression and then tested the effect of “RNA editing events” of these two editing sites on the expression of *LEP*. To evaluate the regulation of *miR-9-5p* on *LEP*, we over-expressed *miR-9-5p* in the HTR-8/SVneo cell line and quantified the expression level of *LEP* using qRT-PCR, and found *LEP* expression decreased significantly compared to control group (**Fig. 4E**), demonstrating that *miR-9-5p* could repress the expression of *LEP*. To test if *LEP* expression was regulated by the editing of the *miR-9-5p* binding sites, we constructed two reporter plasmids carrying a luciferase coding sequence fused to a 3’-UTR of *LEP* which had a *miR-9-5p* targeting region with the RNA editing sites S6 and S7. One plasmid contained wild-type (WT) 3’-UTR with normal S6 and S7 (*LEP*_*WT-UTR*_), the other contained mutated 3’-UTR with A-to-G conversion (A-to-I editing) at S6 and S7 nucleotides (*LEP*_*MUT2-UTR*_). These plasmids were co-transfected with *miR-9-5p* into the HTR-8/SVneo cell line derived from placenta trophoblast, and luciferase activity was examined to test the effect of miRNA on the expression of the reporter gene with edited or not-edited 3’-UTR of *LEP*.

As expected, the luciferase activity was down-regulated by 20% when *miR-9-5p* and *LEP*_*WT-UTR*_ were co-transfected into the cells, compared to transfection of *LEP*_*WT-UTR*_ alone (**Fig. 4F**), demonstrating that the luciferase activity was suppressed by *miR-9-5p* when the luciferase mRNA has wild type 3’-UTR of *LEP*. The luciferase activity was not changed when *miR-9-5p* and *LEP*_*MUT2-UTR*_ are co-transfected, compared to transfection of *LEP*_*MUT2-UTR*_ alone (**Fig. 4F**), suggesting that the luciferase expression was not suppressed by *miR-9-5p* when the luciferase mRNA has mutated 3’-UTR of *LEP*. These results suggest that RNA editing on the *miR-9-5p* binding sites in 3’-UTR may inhibit *LEP* mRNA degradation (**Fig. 4G**).

## Discussion

Genome-scale RNA editing profiling has been done in a series of human normal tissues including brain, lung, kidney, liver, heart and muscle (Blow et al. 2004; Picardi et al. 2015). However, RNA editing has rarely been studied in placenta. Since RNA editing dysregulation is a common phenomenon in cancer, large-scale RNA editing studies on cancer tissues and cell lines have been well documented (Delva et al. 1987; Park et al. 2012; Xu et al. 2018). Recently, 85 genes were found to have differential RNA editing in two placental cell lines treated with hypoxia using transcriptome sequencing (preprint in bioRxiv) (Hasan et al. 2019). Since preeclampsia is thought to be caused by poor placentation, We have recently carried out transcriptome sequencing on placentae from 33 preeclampsia patients (including 9 early-onset severe, 15 late-onset severe and 9 late-onset mild PE patients) and 32 normal subjects, and found a large number of placental dysregulated genes in EOSPE (see related manuscript by Ren et al.). To further find out to what extent the RNA editing might be involved in gene dysregulation in preeclamptic placenta, we performed RNA editome profiling on the placental transcriptomes of the 9 patients with EOSPE and 32 normal controls, and identified a total number of 236,569 RNA editing sites, including 2,920 differentially edited sites (DESs) (**Fig. 1D and 2A**), which can be considered as potential disease-associated sites. The 2,920 DESs include 720 sites in intergenic regions and 2,200 in 1,229 genes, which contains 225 sites in exons, 1,621 in introns and 354 in UTRs (**Fig. 1A and Supplemental Table S4**). DESs in exons, introns or UTRs are enriched with known PE-associated genes respectively (**Fig. 3A**), suggesting that RNA editing sites in both coding and non-coding regions may be involved in PE.

The exonic DESs are mostly C-to-U site-specific changes. Some of these DESs might change the functions of the proteins through generating missense or stop-gain mutations, as reported in many studies on RNA editing in some diseases (Skuse et al. 1996; Anant and Davidson 2001; Mukhopadhyay et al. 2002). Since the majority (73.7%) of the DESs in genes are located in introns, and since the intronic DESs were enriched with DEGs in EOSPE (**Fig. 3B**), we recognized their potential role in the preeclamptic gene dysregulation in the placenta (**Fig. 2A**). RNA editing in the introns affects alternative splicing and RNA degradation in cancer (Kim et al. 2005; Hsiao et al. 2018). The DESs in introns may lead to the gene dysregulation through the same mechanism. RNA editing in UTRs participates in the gene expression homeostasis during cancer development (Yang et al. 2017; Sagredo et al. 2018). Two DESs in the 3’-UTR of *LEP* gene were empirically verified to regulate the gene expression level (**Fig. 4**). We generated “RNA editing events” at miRNA-targeting sites in 3’-UTR of *LEP*, and confirmed that the RNA editing at these sites was able to increased expression level of *LEP* (**Fig. 4**), as observed in patients, suggesting that RNA editing is indeed involved in the gene expression regulation in the placenta of PE patients through modification of miRNA-editing site in 3’-UTRs. RNA editing on miRNA-targeting sites in 3’-UTRs is reported in previous studies in cancer (Yang et al. 2017; Sagredo et al. 2018).

Our comprehensive placental RNA editome dataset contains 236,569 RNA editing sites, 121,749 (51.5%) being placenta specific, which have not been reported in current RNA editome databases, 48.5% being common with other tissues, which have been reported in at least one database. We have identified 2,920 differentially edited sites (DESs) as potential PE-associated RNA editing sites. In conclusion, there is widespread RNA editing in the transcriptome of the placenta, and the dysregulated RNA editing on a group of editing sties may be involved in the development of preeclampsia.

## Methods

### Sample Collection

This research has been approved by the Ethnics Board of Nanfang Hospital, Southern Medical University, China and all patients have signed the informed consent. We collected the tissue samples at the Department of Obstetrics & Gynecology of Nanfang Hospital in China from January 2015 to July 2016. The clinical characteristics of each patient were extracted from the medical records, which strictly followed the American Board of Obstetrics and Gynecology, Williams Obstetrics 24th edition. Preeclampsia (PE) is characterized by new-onset hypertension (≥140/90 mmHg) and proteinuria at ≥20 weeks of gestation, or in the absence of proteinuria, hypertension with evidence of systemic disease (Chaiworapongsa et al. 2014). If patients with PE have systolic blood pressure ≥160 mmHg or diastolic blood pressure ≥110 mmHg, or other organ failures, we call them “preeclampsia with severe features” or briefly “severe PE”. According to gestational age at diagnosis or delivery, PE can be divided as early-onset (< 34 weeks) and late-onset (≥ 34 weeks) (Redman and Sargent 2005). This study focused on the early-onset severe PE (EOSPE). Samples of placental tissues were collected from the mid-section of the placenta between the chorionic and maternal basal surfaces at four different positions, within 5 minutes after delivery. These mid-section placental tissues were placed into RNAlater® solution and stored at −80 °C. The clinical information of the three clinical subtypes and comparisons between subtypes were listed in Supplemental Table S1.

### RNA extraction and RNA-seq

Total RNA was isolated using the RNeasy® Plus Universal Mini Kit (Qiagen, DEU) according to the manual manufacturer’s instruction. Briefly, RNAs with poly(A) tails were isolated, and double-stranded cDNA libraries were prepared using TruSeq RNA Kit (Illumina, USA). The 125-base paired-end sequencing was carried out using Illumina Hiseq 2500 platform (Berry Genomics, CHN).

### Reads alignment and variant calling

Adaptor sequences were trimmed before read alignment. If any read in a pair exhibited more than 4 unidentified bases or more than 63 bases (half of the read length) with base quality score lower than 3, the paired reads would be judged as low quality and discarded. Then RNA-seq reads were indexed using bowtie2 (v2.2.8) and aligned to the genome using tophat2 (v2.1.1) (run with parameters ‘–p6 –r50 –b2 –very–sensitive’) (Langmead and Salzberg 2012; Kim et al. 2013). The reference genome was downloaded from GATK (Genome Analysis Toolkit) (McKenna et al. 2010) hg38 bundles. Duplication levels were evaluated by Picard ‘MakeDuplicates’ (https://github.com/broadinstitute/picard; Broad Institute) (**Supplemental Table S2**). GATK (v3.5) recommended pipeline was modified to suit for RNA-seq data (**Supplemental Materials**). Only variants that passed GATK VQSR (Variant quality score recalibration) filtering were retained. ANNOVAR (v2016-02-01) was used to annotate pass-filtering variants (**Supplemental Materials**) and variants with allele frequency (estimated by 1000 genomes project) smaller than 0.05 in East Asian population (Eas_AF < 0.05) or without allele frequency annotation were selected for further analysis (Wang et al. 2010). Variants with ANNOVAR annotated as ‘splicing’, ‘upstream’ or ‘downstream’ would be collapsed into ‘intergenic’ category.

### Detection of potential RNA editing sites

Considering that our sequencing library was not strand specific, we extracted the variants with A/G and C/T mismatch or T/C and G/A mismatch corresponding to plus and minus strand. These detected variants were taken as potential as A-to-I and C-to-U RNA editing sites.

### Validation of candidate RNA editing sites

We collected 2,576,459 known RNA editing sites from database RADAR (version 2) (Ramaswami and Li 2014), 314,250 from DARNED (Kiran and Baranov 2010) and 4,668,508 from REDIportal (Picardi et al. 2017). Since these RNA editing databases were annotated using hg19 version genome, we converted coordinates of the RNA editing sites we detected to corresponding position in hg19 genome through LiftOver tool in UCSC (http://genome.ucsc.edu/cgi-bin/hgLiftOver) and 235,488 sites were successfully converted. The editing sites detected at mRNA level were re-evaluated by multiple sequence alignment of reads that covered the RNA editing sites.

For empirical validation, we randomly chosen 33 regions with 89 RNA editing sites, including 17 sites from exons, 37 from UTRs and 35 from introns (**Supplemental Table S3**). The regions with RNA editing sites were PCR-amplified from gDNA from placenta samples (9 EOSPE vs. 9 normal samples; Supporting Information) and then Sanger-sequenced for validation of the editing events (Sangon Biotech, CHN). Sites without gDNA mutations at the corresponding potential RNA editing sites (A-to-G or C-to-T on mRNA) were considered as validated RNA editing sites.

### Detection of EOSPE differentially edited sites (DESs)

We evaluated the RNA editing level for a given site by RNA editing ratio, which is defined as the ratio of reads with editing to total reads. If there is no read covering the editing site in a sample, editing ratio for this site in this sample would be assigned as zero. To detect reliable differentially edited sites (DESs) in placenta, we set filtering criteria as: (i) having no less than three reads that supported the editing site and no less than five reads that covering the editing site in at least three samples; (ii) no more than three (∼30%) EOSPE samples that had editing ratio = 1; (iii) no more than nine (∼30%) normal samples that had editing ratio = 1. A total number of 23,645 editing sites were kept for DES detection. After classified editing sites into EOSPE up-editing and EOSPE down-editing groups, we compared the editing ratios between EOSPE and normal placentae by one-tailed Wilcoxon test for these two groups. Then we selected RNA editing sites exhibited significantly editing ratio differences between EOSPE and normal placentae (*P <* 0.05) as DESs.

### Detection of EOSPE differentially expressed genes (DEGs)

We mapped clean reads to human genome (hg38 version) by STAR (v2.5.2b) and called raw counts for every gene in every sample by HTSeq (v0.9.0) htseq-count separately (Dobin et al. 2013; Anders et al. 2015). Differentially expressed genes (DEGs) were detected using DESeq2 (v1.16.1) (Love et al. 2014) and edgeR (v3.18.1) (Robinson et al. 2010), and intersection of results from these two detection methods were subsequently taken as final DEGs (2,977 genes). Human reference genome and gene annotations were downloaded from Ensembl.

### Collection of known PE-associated genes

Known PE-associated genes were collected from eight published papers. After dataset merging, totally 1,177 genes were referred as PE-associated genes for further analysis (**Supplemental Table S5**).

### Enrichment calculation of EOSPE DEGs and PE-associated genes in genes with DESs

According to the enrichment calculation of hypergeometric distribution, we calculated enrichments for DEGs in EOSPE and PE-associated genes by comparing the fraction of these genes in foreground and background dataset. Genes with DESs and all expressed genes in EOSPE and normal placentae were referred as foreground and background dataset. A gene was defined as expressed gene if there is at least one read in all samples of both EOSPE and normal placentae. One tailed Fisher-exact test was used to calculate the *P*-value. The standard error of the fraction was estimated using a bootstrapping method with 10,000 resamplings (randomly extracted 10% genes from foreground and background per resampling).

### Functional enrichment of GO terms and KEGG pathways in genes with DESs

Functional enrichment analysis was achieved by ClulsterProfiler (v3.10.1) through R (Yu et al. 2012). Threshold of adjusted *P*-values were set to 0.05 to find significantly enriched GO terms or KEGG pathways.

### Association of differential RNA editing with differential gene expression

For every differentially edited site (DES), we re-classified all 41 samples into two groups: edited group and not-edited group based on the existence of the editing event. DESs with less than three samples in edited or not-edited group were removed. Genes without expression in any of the samples were also removed. We then compared the gene expression between edited group and not-edited group by one-tailed Wilcoxon test. The DESs which had significant gene expression differences between edited group and not-edited group (*P <* 0.05) were considered as had association with the differential gene expression. Counts per million bases calculated by edgeR were used as gene expression here.

### Correlation between differential RNA editing and gene expression level

For a given DES, we calculated the correlation between editing ratio and gene expression level by Hmisc (https://cran.r-project.org/web/packages/Hmisc/index.html; v4.2-0) in R. Genes without expression in any of the samples were removed. Counts per million bases calculated by edgeR were used as gene expression. *P =* 0.05 was set as cutoff for significant correlation.

### Protein-protein interaction (PPI) network of *RHOA* and EOSPE DEGs

Human interactome data was from InWeb_InBioMap (https://www.intomics.com/inbio/map) and *RHOA*-centered PPI network was extracted from human interactome network containing *RHOA* and its interactors (Li et al. 2017). Then EOSPE DEGs were mapped onto *RHOA*-centered network and *RHOA*-subnetwork of DEGs in EOSPE was extracted. The network extraction was done using CytoScape (v3.4.0) (Shannon et al. 2003).

### Identifying intron retention (IR) event

We downloaded GENCODE canonical transcript annotation (v29) from UCSC Table Browser (Karolchik et al. 2004). The transcripts mapped to scaffolds were removed. If a gene had multiple transcripts mapped to different chromosomes, the gene would also be removed. Then we got a set of genes with one or more canonical transcripts. If the gene had more than one canonical transcripts, we merged them into one transcript by keeping the nonoverlapping exons and collapsing the overlapping exons. The final set of canonical transcripts were unique to their corresponding genes.

We assembled transcripts from RNA-seq reads by StringTie (v1.3.4) (Pertea et al. 2015), and compared them with known transcripts (ftp://ftp.ensembl.org/pub/release-84/gtf/homo_sapiens/Homo_sapiens.GRCh38.84.gtf.gz) by cuffcompare in Cufflinks (v2.2.1) (Trapnell et al. 2010). We further compared the assembled transcripts with corresponding final set of canonical transcripts, and identified the regions of the assembled transcripts that fell into the introns of the canonical transcripts. These regions were defined as IR events.

### Calculation of percent spliced in (PSI)

For every IR event, we calculated the percent spliced in (PSI) according to the below formula (Schafer et al. 2015): 

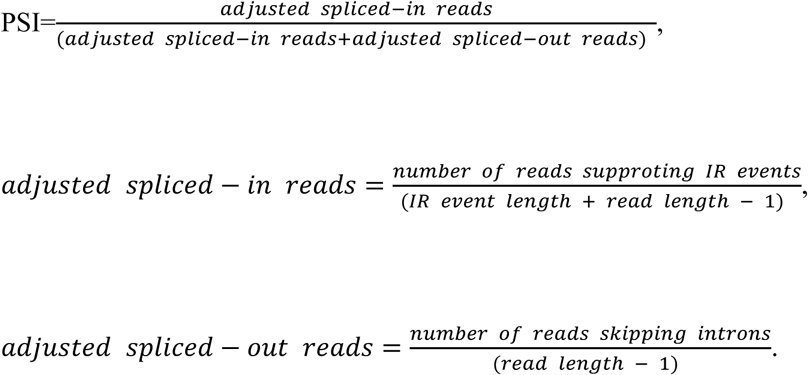

### Association of intronic differential RNA editing with intron retention

We compared the percent spliced in (PSI) of each intron retention between edited group and not-edited group by one-tailed Wilcoxon test. For a DES, if RNA editing happened in less than three samples within either edited group or non-editing group, it was removed. The DESs which had significant PSI differences between edited group and not-edited group (*P <* 0.05) were considered as associated with the IR events.

### Prediction of miRNA-targeting genes harboring DESs

Human miRNA-targeting information was downloaded from TargetScan database (v7.2) (Shi et al. 2017), and used for identifying DESs in miRNA-targeting regions.

### Estimation of relative expression level of LEP and miR-9-5p in placentae

For *LEP* and *miR-9-5p*, the mRNA expression level was evaluated by quantitative real-time quantitative PCR (qRT-PCR) in 8 EOSPE samples and 8 randomly-chosen normal samples. *ACTB* and U6 were used as internal control for *LEP* and *miR-9-5p* respectively. Expression levels of mRNAs were normalized to internal control mRNA. The relative expression levels of the genes were calculated using the 2-ΔΔCt method. ΔCt was the difference of Ct values between quantification of *LEP* and *GAPDH* or between *miR-9-5p* and U6. The standard error of 2-ΔΔCt values was calculated to serve as error bar. The primers used for mRNA quantification were listed in Supporting Information. Wilcoxon test was used to calculate the significance of the expression difference between EOSPE and normal group using R package.

### Cell culture

HTR-8/SVneo cell line was purchased from ATCC (USA). Cells were grown in RPMI 1640 media (Gibco, Thermo Fisher Scientific, USA) supplemented with 10% (v/v) fetal bovine serum (FBS) and incubated at 37 °C and 5% (v/v) CO_2_.

### Quantitative real-time PCR (qRT-PCR) experiment

The *miR-9-5p* mimic or negative control (NC) mimic (Shanghai GenePharma Co., Ltd, CHN) was transfected into HTR-8/SVneo cell line by Lipofectamine 2000 (Invitrogen, Thermo Fisher Scientific, USA) in six-well plates, and cells were harvested after 48 hours of transfection. RNAs were extracted using RNAeasy Mini Kit (Qiagen, USA) according to the manufacturer’s instruction. The RNAs were reverse - transcribed into cDNAs using the PrimeScriptTM RT Kit (Takara, JPN) following the manufacturer’s instruction. *ACTB* was used as the internal control. The qRT-PCR of *LEP* mRNA quantification in HTR-8/SVneo cell line was the same as described above. The relative expression levels of the genes were calculated using the 2-ΔΔCt method. Wilcoxon test was used to calculate the significance of the expression difference between the cells transfected with *miR-9-5p* mimic and with negative control mimic using R package. The standard error of 2-ΔΔCt values was calculated to serve as error bar.

### Luciferase activity experiment

Two luciferase-reporter plasmids were constructed, one carrying non-edited *miR-9-5p* target region (*LEP*_*WT-UTR*_), another carrying S6 and S7 edited *miR-9-5p* target region of *LEP* (*LEP*_*MUT2-UTR*_). The day before transfection, HTR-8/SVneo cells were plated at 1 × 5,000 cells per well in 96-well plates. Then reporter plasmids were co-transfected with *miR-9-5p* mimic or negative control (NC) into HTR-8/SVneo cell line and cultured in 96-well plates using TurboFect (Thermo Fisher Scientific, USA) according to the manufacturer’s instruction. In each well, the cells were transfected with 100 ng of the tested constructs and 20 ng of pRL-TK. The activities of Renilla luciferase and Firefly luciferase were determined using the Dual-Luciferase Reporter Assay System (Promega, USA) 48 hours after transfection. Luminescence was detected by a GloMax 20/20 Luminometer (Promega, USA) according to the manufacturer’s instructions. The error bars represent standard errors.

## Supporting information

Supplemental Figure 2, panel A and B

Supplemental Figure 2, panel C

## Data Access

The authors are committed to the release of data and analysis results, with the understanding that rapid and transparent data sharing will benefit the scientific research community. Detailed analyzed data are included in supplementary excel tables. RNA-seq data are going to be deposited at GEO and will be available. All renewable reagents and detailed protocols will be available upon request.

## Acknowledgements

The work was supported by the National Natural Science Foundation of China grant No. 81571097 (X.Y.) and grant No. 81671466 (M.Z.), the Natural Science Foundation of Guangdong Province Grant No. 2016A030308020 (X.Y.), Science and Technology Projects of Guangzhou grant No. 201704020116 (X.Y.), and the Science and Technology Planning Project of Guangdong Province grant No. 2015B050501006 (M.Z.). The funding organizations had no role in design and conduct of the study; collection, management, analysis, and interpretation of the data; and preparation, review, or approval of the manuscript.

## Author contributions

Xinping Y. conceived the project; Xinping Y., M.Z. and Y.Y. supervised the project. Xiaoxue Y. performed computational analysis with contributions from H.L., Y.G., and Z.R.; J.Y., X.L. Q.C., S.J. performed the experiments. Xiaoxue Y. and Xinping Y. wrote the paper.

## Disclosure Declaration

The authors declare no competing interests.

## Notes

#### Summary of Updates

Added supplementary data and materials.

