## Supplemental Figure 1 for "Landscape of Dysregulated Placental RNA Editing Associated with Preeclampsia"

### Supplementary Figures

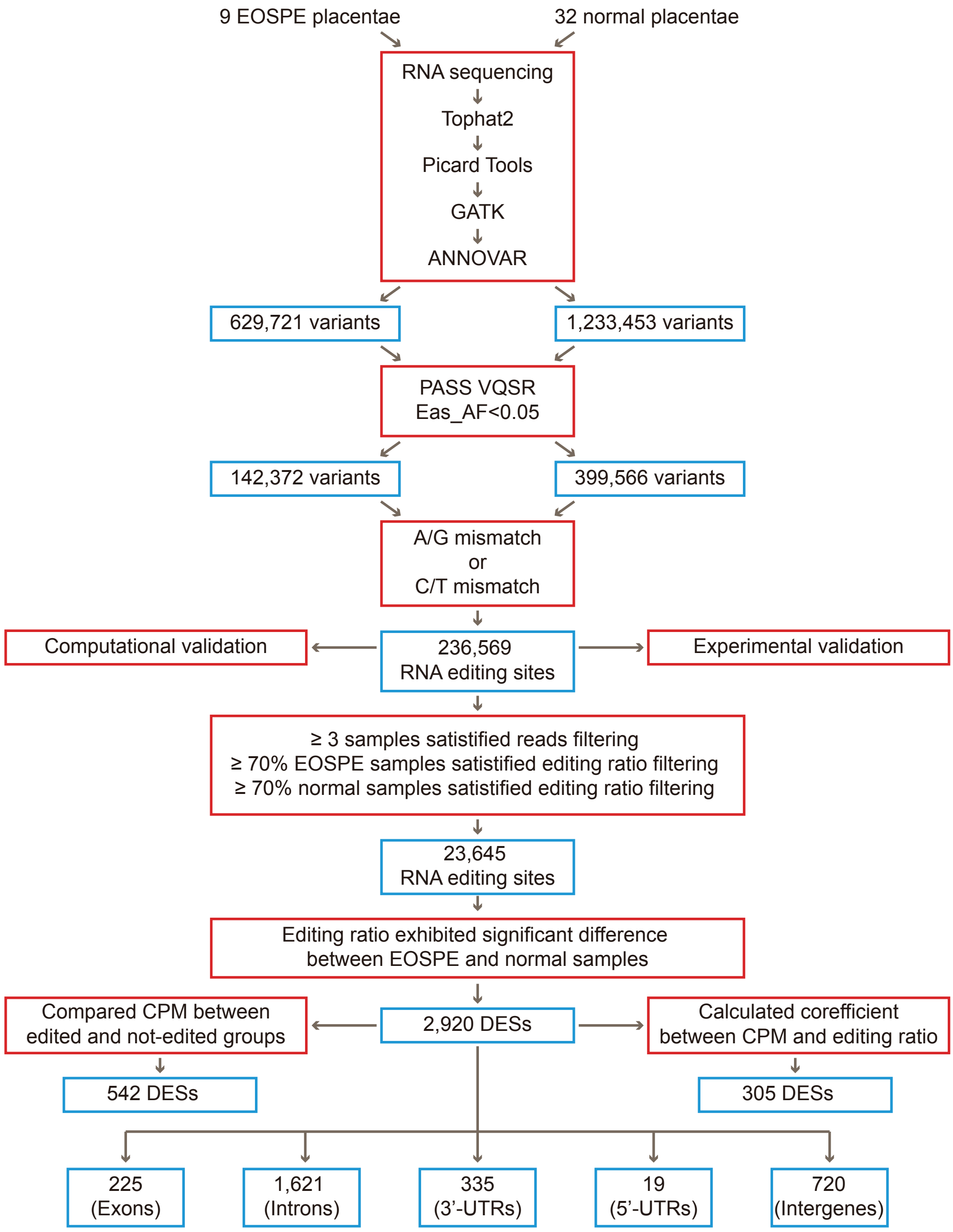


**Supplementary Figure S1.** Identification, detection and analysis of RNA editing sites in placentae. Red boxes represent the analysis strategies or filtering criteria. Blue boxes represent the results. VQSR: variant quality score recalibration; Eas_AF: allele frequency in East Asian population; DES: differentially edited site; CPM: counts per million bases; UTR: untranslated region
