## Supplementary figures and images for "Landscape of Dysregulated Placental RNA Editing Associated with Preeclampsia"

### Supplemental Figure 2, panel A and B

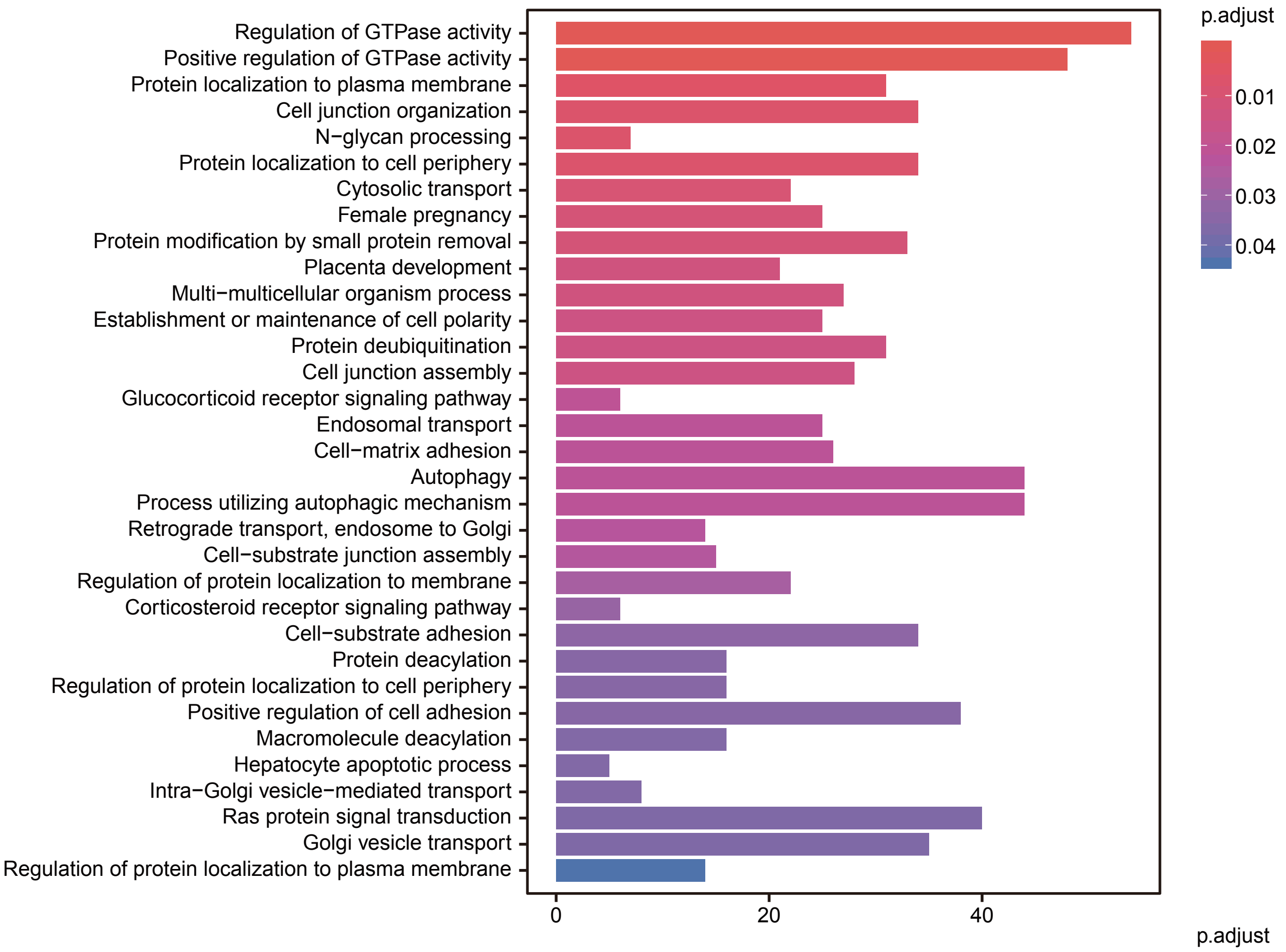

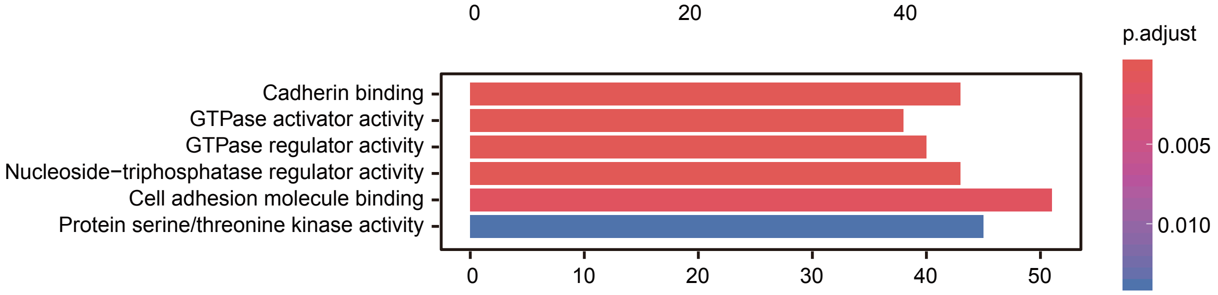
