## Supplemental Figure 2, panel C for "Landscape of Dysregulated Placental RNA Editing Associated with Preeclampsia"

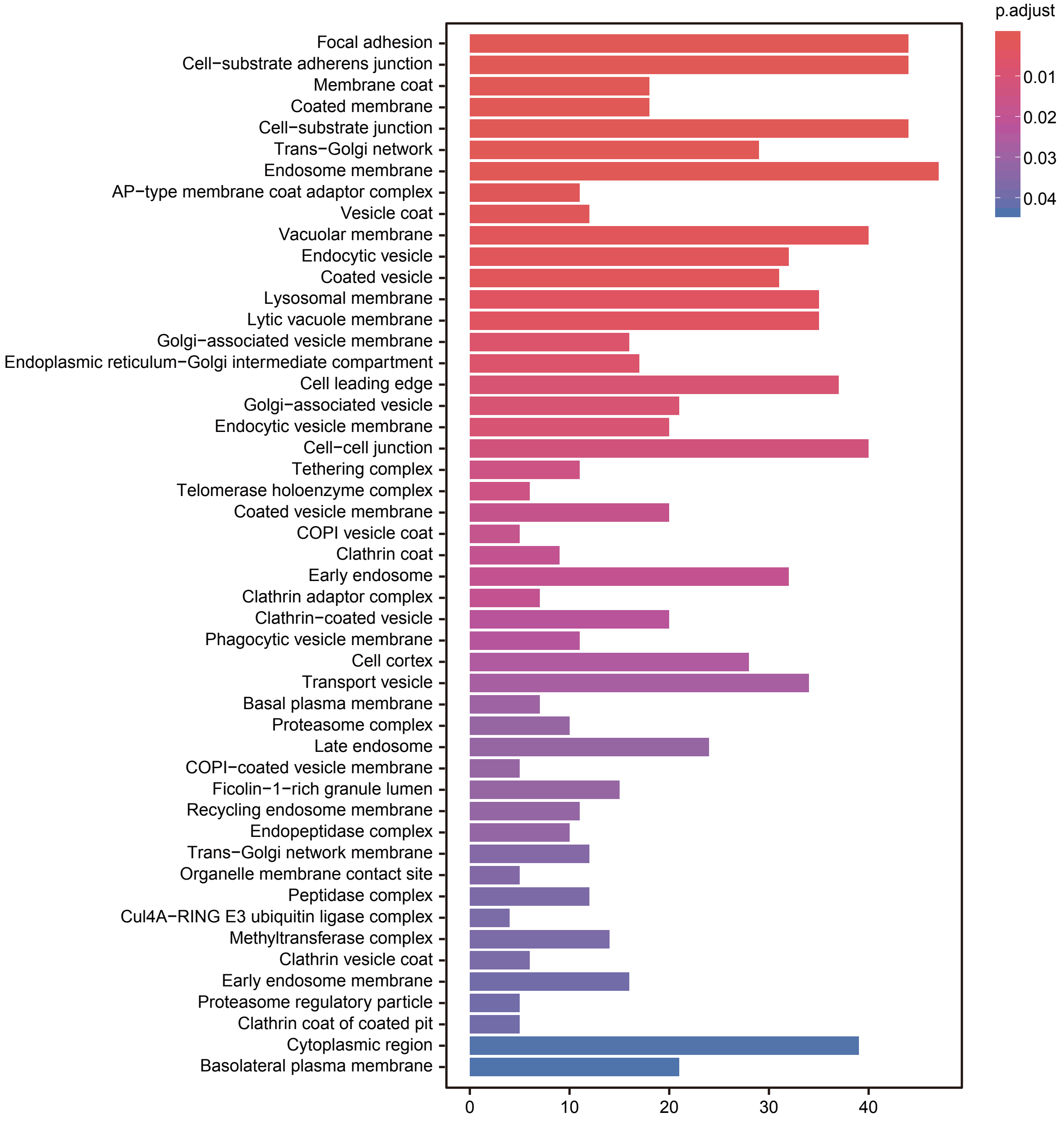


**Supplementary Figure S2.** Enrichment of genes with DESs in GO terms. (A) The GO BP terms enriched with genes harboring DESs. (B) The GO MF terms enriched with genes harboring DESs. (C) The GO CC terms enriched with genes harboring DESs. Color of bars: adjusted *P*-values of the significance of the enrichments; X-axis: the number of genes with DESs in the terms. Enrichment analysis was carried out using ClusterProfiler.
