## Supplemental Figure 3 for "Landscape of Dysregulated Placental RNA Editing Associated with Preeclampsia"

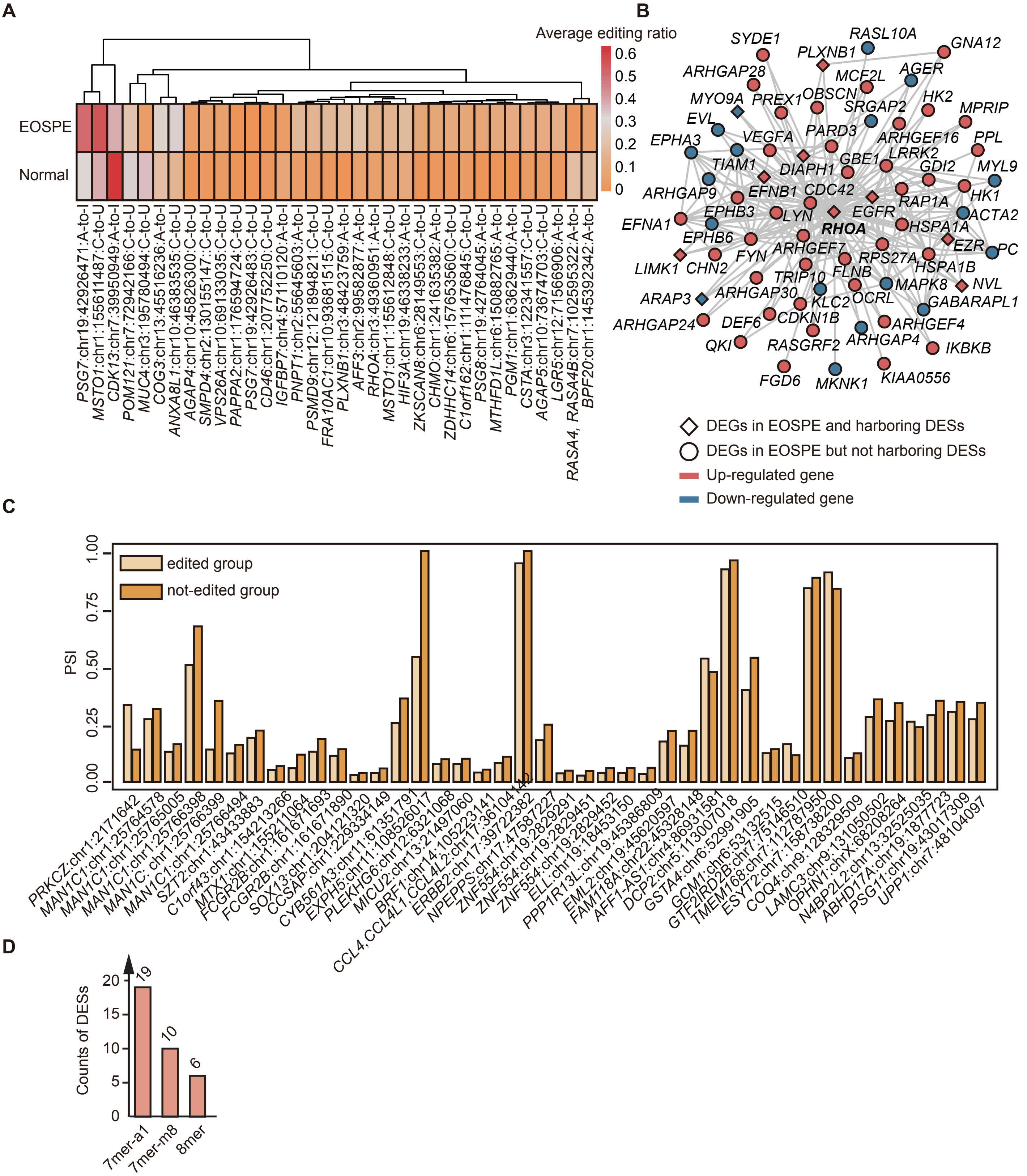


**Supplementary Figure S3.** DESs in exons, introns and 3'-UTRs. (A) Heatmap of average editing ratio of exonic nonsynonymous/stopgain DESs. Values of average editing ratio were indicated by colored bars. (B) Protein-protein interaction networks of gene *RHOA*. All nodes in the network are DEGs of EOSPE, some harboring DESs. Diamonds and circles represent the DEGs with/without DESs respectively. Red represents up-regulation and blue down-regulation in EOSPE. (C) Intron retention (IR) events with significant difference of PSI (percent spliced in) between the edited group and not-edited group. Samples were classified into edited group or not-edited group based on the existence of RNA editing at a given site. X-axis represents the IR events with intronic DESs. Y-axis represents the means of PSI in edited group or not-edited group for a given DES. (D) Seed match types of miRNA-targeting regions in 3'-UTR harboring DESs. The miRNA-targeting regions and the seed match types were predicted using TargetS-can. MiRNA-targeting regions with null PCT (preferentially conserved targeting) values were removed. Numbers above the bars are the number of DESs.
