## Supplemental Data for "Landscape of Dysregulated Placental RNA Editing Associated with Preeclampsia"

### Placental Editome Profiling Reveals Widespread RNA-Editing Dysregulation in Preeclampsia

**Table of Contents**

Supplementary Figures: page 3-6
Supplementary Tables: page 7-8
Supplementary Materials and Methods: page 9-11

#### Supplementary Figures

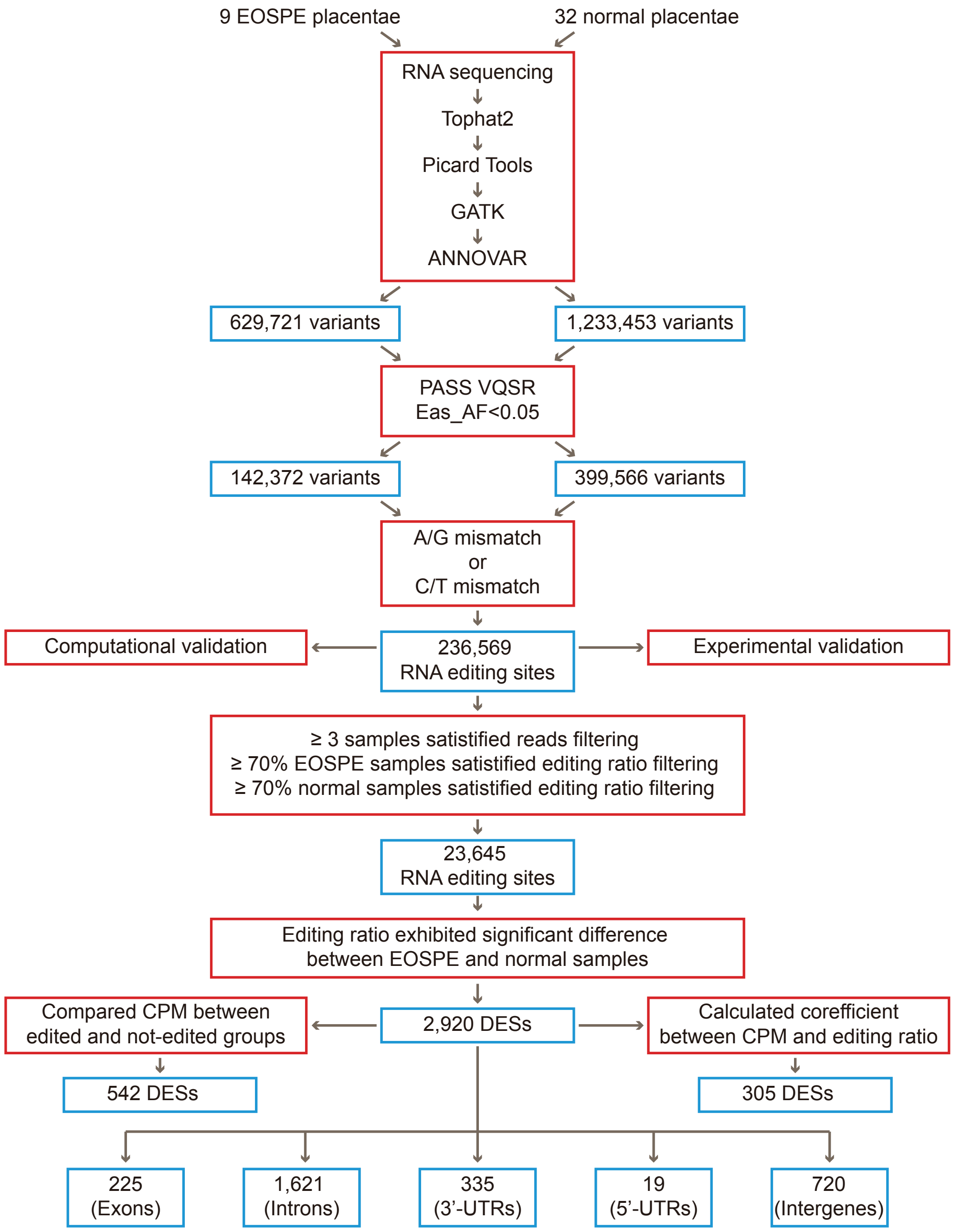


**Supplementary Figure S1.** Identification, detection and analysis of RNA editing sites in placentae. Red boxes represent the analysis strategies or filtering criteria. Blue boxes represent the results. VQSR: variant quality score recalibration; Eas_AF: allele frequency in East Asian population; DES: differentially edited site; CPM: counts per million bases; UTR: untranslated region

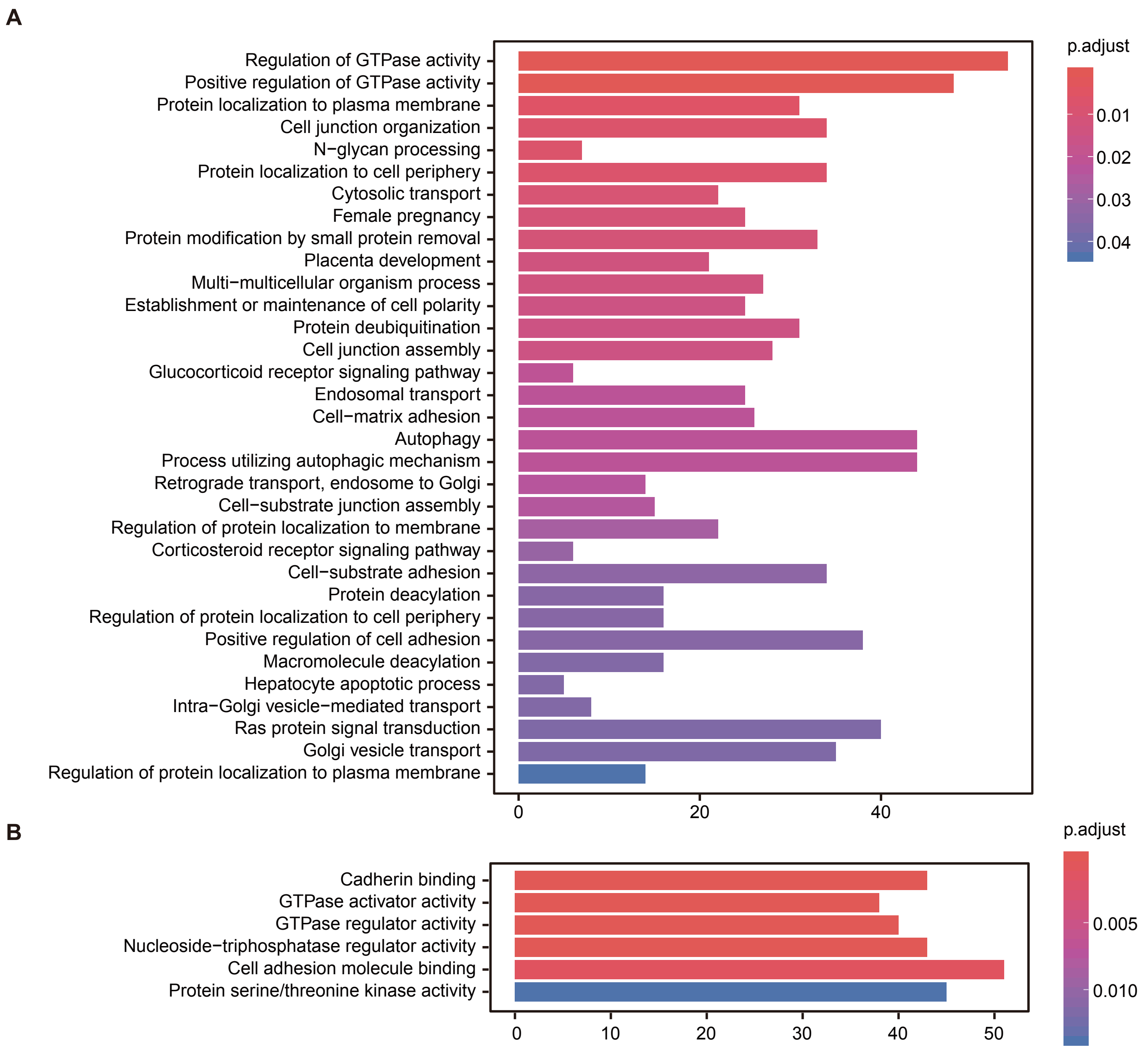
A
Regulation of GTPase activity
Positive regulation of GTPase activity
Protein localization to plasma membrane
Cell junction organization
N−glycan processing
Protein localization to cell periphery
Cytosolic transport
Female pregnancy
Protein modification by small protein removal
Placenta development
Multi−multicellular organism process
Establishment or maintenance of cell polarity
Protein deubiquitination
Cell junction assembly
Glucocorticoid receptor signaling pathway
Endosomal transport
Cell−matrix adhesion
Autophagy
Process utilizing autophagic mechanism
Retrograde transport, endosome to Golgi
Cell−substrate junction assembly
Regulation of protein localization to membrane
Corticosteroid receptor signaling pathway
Cell−substrate adhesion
Protein deacylation
Regulation of protein localization to cell periphery
Positive regulation of cell adhesion
Macromolecule deacylation
Hepatocyte apoptotic process
Intra−Golgi vesicle−mediated transport
Ras protein signal transduction
Golgi vesicle transport
Regulation of protein localization to plasma membrane
0
20
40
p.adjust
0.01
0.02
0.03
0.04
B
Cadherin binding
GTPase activator activity
GTPase regulator activity
Nucleoside−triphosphatase regulator activity
Cell adhesion molecule binding
Protein serine/threonine kinase activity
0
10
20
30
40
50
p.adjust
0.005
0.010

C

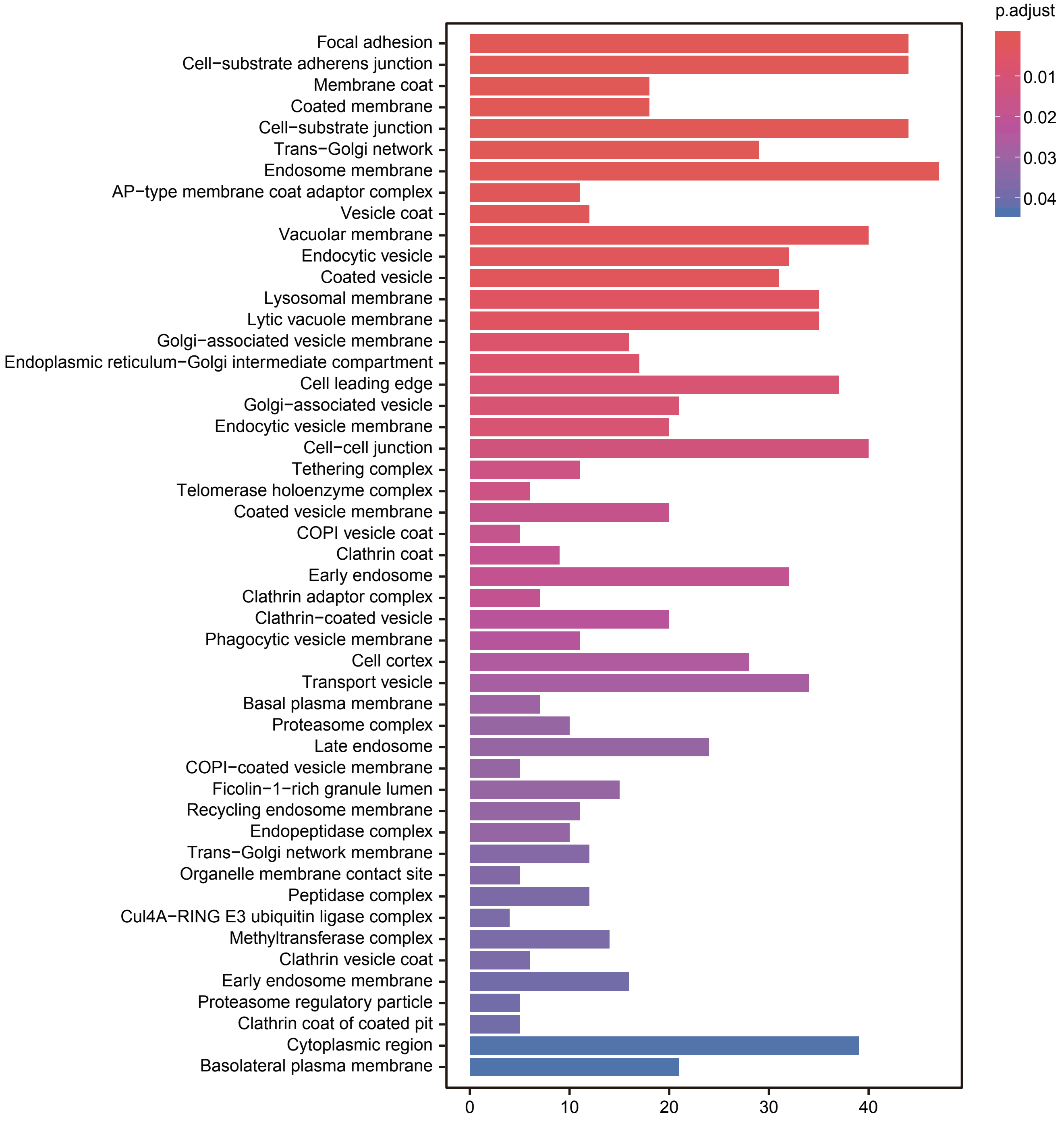


**Supplementary Figure S2.** Enrichment of genes with DESs in GO terms. (A) The GO BP terms enriched with genes harboring DESs. (B) The GO MF terms enriched with genes harboring DESs. (C) The GO CC terms enriched with genes harboring DESs. Color of bars: adjusted *P*-values of the significance of the enrichments; X-axis: the number of genes with DESs in the terms. Enrichment analysis was carried out using ClusterProfiler.

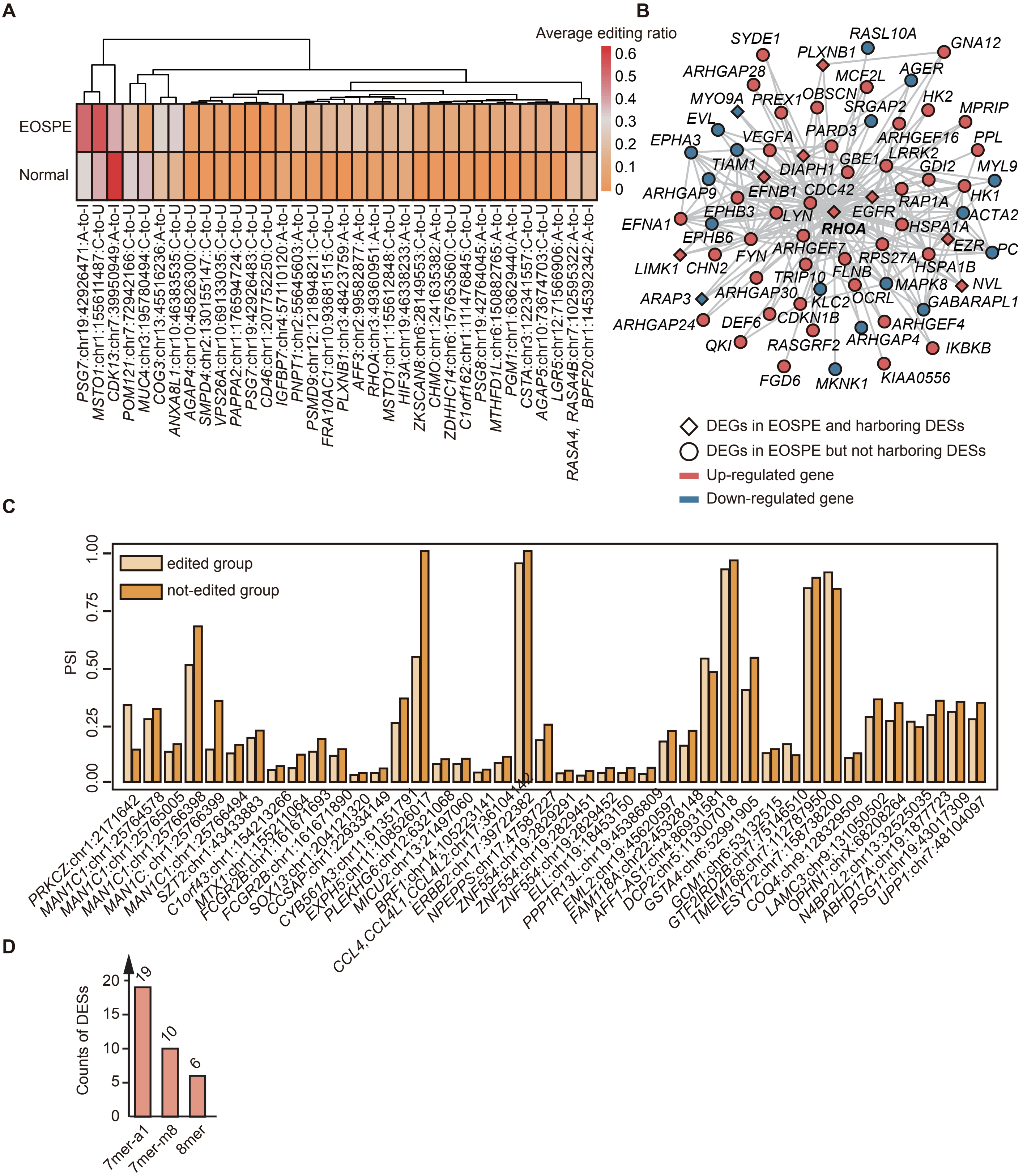


**Supplementary Figure S3.** DESs in exons, introns and 3'-UTRs. (A) Heatmap of average editing ratio of exonic nonsynonymous/stopgain DESs. Values of average editing ratio were indicated by colored bars. (B) Protein-protein interaction networks of gene *RHOA*. All nodes in the network are DEGs of EOSPE, some harboring DESs. Diamonds and circles represent the DEGs with/without DESs respectively. Red represents up-regulation and blue down-regulation in EOSPE. (C) Intron retention (IR) events with significant difference of PSI (percent spliced in) between the edited group and not-edited group. Samples were classified into edited group or not-edited group based on the existence of RNA editing at a given site. X-axis represents the IR events with intronic DESs. Y-axis represents the means of PSI in edited group or not-edited group for a given DES. (D) Seed match types of miRNA-targeting regions in 3'-UTR harboring DESs. The miRNA-targeting regions and the seed match types were predicted using TargetScan. MiRNA-targeting regions with null PCT (preferentially conserved targeting) values were removed. Numbers above the bars are the number of DESs.

**Supplementary Tables S1~S10**

**Supplementary Table S1.** Clinical information of patients with EOSPE and normal subjects. Outlier values in the group assigned with '*' were removed. P-values were calculated using t-test.

| **Clinical information** | **Normal** | **EOSPE** | ***P*-value** |
|---|---|---|---|
| Individual | 30 | 9 | -- |
| Placenta | 32 | 9 | -- |
| Baby | 35 | 10 | -- |
| Age (Year) | 30.57 ± 5.12 | 31.89 ± 6.58 | 5.90E-01 |
| BMI (Kg/m^2) | 28.20 ± 2.78 | 26.62 ± 2.14 | 1.06E-01 |
| Systolic pressure (mmHg)* | 118.70 ± 10.41 | 181.40 ± 17.73 | **1.84E-16** |
| Diastolic pressure (mmHg)* | 73.43 ± 9.63 | 115.60 ± 14.47 | **3.24E-05** |
| Gestational age of delivery (Day) | 273.90 ± 8.45 | 216.80 ± 15.20 | **1.18E-06** |
| Baby weight (Kg) | 3.15 ± 0.55 | 1.35 ± 0.48 | **1.69E-08** |
| Low birth weight (LBW) | 5 (14.3%) | 9 (90%) | **1.98E-05** |
| Proteinuria (g/24h)* | -- | 9.20 ± 1.90 | -- |
| C-reactive protein level* | -- | 2.66 ± 1.44 | -- |

**Supplementary Table S2.** Reads alignment and duplicates evaluation. Parameters were extracted from evaluation results of tophat2 and picard 'MarkDuplicates' program. Available as separate xlsx-file.

**Supplementary Table S3.** Validation of 89 RNA editing sites by sanger sequencing and multiple alignment of the reads. Since mRNA of the sample PHNPR35 ran out during the validation, we used the mRNA of PHNPR34 for validation for 10 regions. Available as separate xlsx-file.

**Supplementary Table S4.** Differentially edited sites (DESs) of EOSPE. Sheet1: DESs with higher editing ratio in EOSPE placentae; Sheet2: DESs with lower editing ratio in EOSPE placentae. Available as separate xlsx-file.

**Supplementary Table S5**. Preeclampsia-associated (PE-associated) genes collected from literature. Sheet1: Literature information, numbers of PE-associated genes and datasets used in these studies; Sheet2: PE-associated genes with times found in the literatures. Available as separate xlsx-file.

**Supplementary Table S6.** Differentially expressed genes (DEGs) of EOSPE. Available as separate xlsx-file.

**Supplementary Table S7**. Enrichment of genes with DESs in GO terms and KEGG pathways. Sheet1: The enriched GO biological process (BP) terms for genes with DESs; Sheet2: The enriched GO molecular function (MC) terms for genes with DESs; Sheet3: The enriched GO cellular component (CC) terms for genes with DESs; Sheet4: The enriched KEGG pathways for genes with DESs. Q-values smaller than 0.05 would be identified as significantly enriched terms/pathways. Available as separate xlsx-file.

**Supplementary Table S8**. Association of DESs with differential gene expression and correlation of RNA editing ratio with gene expression level. Available as separate xlsx-file.

**Supplementary Table S9**. DESs in 3'-UTRs that located in miRNA target regions. Available as separate xlsx-file.

**Supplementary Table S10**. qRT-PCR and luciferase assay results. Available as separate xlsx-file.

**Supplementary Materials and Methods**

**Transcriptome variant calling using modified GATK pipeline**

GATK has recommended variant calling pipelines for whole genome sequencing or exome sequencing data. We have modified analysis pipeline for RNA-seq dataset. For the analysis of the sequencing data of every sample, the pipeline started from bam files after duplicate reads marking by Picard Toolkit "MarkDuplicates" followed by following steps to identify variants:

- SplitNCigarReads: with parameters '*–U ALLOW_N_CIGAR_READS –fixNDN*'.
- RealignerTargetCreator: used reference dataset '*known_indels.vcf*' and '*gold_standard.indels.vcf*'
- IndelRealigner: with parameter '*–U ALLOW_N_CIGAR_READS*' and reference dataset '*known_indels.vcf*' and '*gold_standard.indels.vcf*'.
- BaseRecalibrator: with parameter '*–U ALLOW_N_CIGAR_READS*' and reference dataset '*known_indels.vcf*', '*gold_standard.indels.vcf*' and '*dbsnp138.vcf*'
- PrintReads: with parameter '*–U ALLOW_N_CIGAR_READS*'
- HaplotypeCaller: with parameters '*-dontUseSoftClippedBases –U ALLOW_N_CIGAR_READS –stand_call_conf 20.0 –stand_emit_conf 20.0 –ERC GVCF –nda*'. Since it would require too much computational memory for calling variants by HaplotypeCaller with all chromosomes at one time, we ran HaplotypeCaller for every chromosome separately.
- CatVariants: with parameter '*-assumeSorted*'. This step was aim to combine single chromosome results into one result for every sample.

The variants identified in 41 samples were joined into two files according to sample group, followed by the following steps for "variant quality score recalibration" (VQSR):

- GenotypeGVCFs
- VariantAnnotator: with parameters '*-U ALLOW_N_CIGAR_READS –A Coverage –A QualByDepth –A FisherStrand –A StrandOddsRatio –A MappingQualityRankSumTest –A ReadPosRankSumTest –A RMSMappingQuality*'. This step was aim to add additional annotations required by VariantRecalibrator.
- VariantRecalibrator: with parameters '*-an QD –an MQ –an MQRankSum –an ReadPosRankSum –an FS –an SOR –an InbreedingCoeff –mode SNP –tranche 100 –tranche 99.9 –tranche 99 –tranche 97 –tranche 95 –tranche 93 –tranche 90 –resource:hapmap,known=false,training=true,truth=true,prior=15.0 –resource:omni,known=false,training=true,truth=true,prior=12.0 –resource:1000G,known=false,training=true,truth=false,prior=10.0 –resource:dbsnp,known=true,training=false,truth=false,prior=2.0*' and reference dataset '*hapmap_3.3.vcf*', '*1000G_omni2.5.vcf*', '*1000G_phase1.snps.high_confidence.vcf*' and '*dbsnp138.vcf*'.
- ApplyRecalibration: with parameters '*-mode SNP –ts_filter_level 99.5*'.

All reference datasets used in GATK analysis were included in GATK hg38 version resource bundle.

**Variant annotations using ANNOVAR**

After filtered based on VQSR results, the variants were further annotated using ANNOVAR, which was a recommended software in GATK guidebook. Reference files for annotation were downloaded through ANNOVAR 'annotate_variation.pl' program with parameters '*-buildver hg38 –downdb' or '-buildver hg38 –downdb –webfrom annovar*'. Details were listed below:

- *refGene:* v20170601.
- *cytoBand:* downloaded at 20140511.
- *esp6500siv2_all:* v20141222.
- *1000g2015aug:* including *1000g2015aug_all*, *1000g2015aug_afr*, *1000g2015aug_eas* and *1000g2015aug_eur;* v20150824.
- *avsnp147:* v20160606.
- *dbnsfp30a:* v20170221.

**PCR primers for the validation of RNA editing sites**

To amply PCR for RNA editing validation, we randomly chosen 33 regions including 89 RNA editing sites. Then PCR primers were designed to include about 400 bp covering the picked RNA editing sites.

| No | Forward PCR primer (5' ~ 3') | Reverse PCR primer (5' ~ 3') |
|---|---|---|
| 1 | TCTTGTCTCAATACTGCTTTGG | GTATGCCACCACATCTACCT |
| 2 | TGAAGCCTAAACCAAACCAG | GACCAGAAGTTCAAGACCAG |
| 3 | ACCATGACTGCAAAGTTCCT | TTTAATCTCCAAGCTCCACTCC |
| 4 | CTTCTTATTAGCTGGACGACCT | CTTACTAAGTCCCAGGCACC |
| 5 | CCAGACCACAATCTTCAAAGAG | TAATAGAAGACTGGGTGCGG |
| 6 | GCTAGAAGATACATGTCCACAC | TTCTCGAAGACCTCACATCAC |
| 7 | GACAAACATACAGAAATGCACC | CATGGATTCAAGGCTGTTGTG |
| 8 | GGAAATTGAGGCCACAAGAG | CCTTCCACTCAGCTTTATTCAC |
| 9 | CACAGATGCCAATGAGTCAG | CATGTGATTCATACAGTCCGT |
| 10 | TGACACAGTGGTTCTCAACCG | TGACATGCATGGTCCTCATAGTG |
| 11 | TGGCTCACGCTTATAGTTCC | GTGTCTCAAATTATCCCATACGTC |
| 12 | ACTTGCTGAGGGTGAAGTCC | TCCTGGTCAAAGAGGCTTGG |
| 13 | GGATTTAGAGTCTGAATGGCAGC | GACCAGACACAAGGCAATACTG |
| 14 | GCAATACTGGAACTCACCAC | CAAGCTGCCTTCTCCTATGAG |
| 15 | AGAGTAGAGCCATAGAAAGGCA | GGTTTCAAGGCATTGAGGAC |
| 16 | AATATCTGAGTCACAGTGTCCA | ATAAACAGTCAGCAAATGGCAG |
| 17 | CCCTTAATGGCCAGCCTGGATG | CCAGGAGCACTTACTCATTGAAAAC |
| 18 | GGAGGAAGGCAAGGGTATTTTTTAC | GGTGGGTAAAGAGACACAGGGC |
| 19 | ATCATTGCAGACAAAACTACCAG | ATTACAGGCACCTGCCACC |
| 20 | CTCTTGTCTCCCAGGCTGG | CCTGCTCAAGACCGGACAC |
| 21 | CTCTGTTGTCCAGGCTGGAG | GAAGGAGCCTCGGCCAGG |

| | | |
|---|---|---|
| 22 | ACCGAGGCTCAGGGGTATC | TCACCTTCACAGAGAAGGGGC |
| 23 | ATTCTGTACAGTTTCAGGTTCTTC | GAGGCTGGACAGCCACAG |
| 24 | CCTCCTGAGTAGCTGGGAC | AAGCTAAACAAGGAGGCATCAG |
| 25 | GGGAGGTTGAGGATGATGCAG | GCCCAGCTCAAAGTGAAAGTG |
| 26 | GCCCCCGTCACTGTGGGTG | GGGAGTCGGGGGAAGGGAG |
| 27 | GTGAGGTCTTTTGCTACAAGGAGG | CCCCACAAGCGGAATTGTATTG |
| 28 | GAGTGCAGTGGCGCAGTC | GTCCGGCTGGGAAAACCC |
| 29 | CACACCCTTAACCTTTAAGCCAG | ATTAGCCGGGCGTGATGG |
| 30 | AGATTCGAACACAGTGACTCAAGG | GGGCTGGGGAGTCATGGTG |
| 31 | CCCTGTGCTTCGTCCACAG | CCACGTTCCCTGCTGCCTC |
| 32 | CCATGTGGATAAAATGCCAGAC | GACCAGCCTAGCCAATATGGTG |
| 33 | TCTCCGGTGGCCCGGCCC | CCGGTGCCCCCAAGGAGAG |

**Primers for qRT-PCR of *LEP* and *miR-9-5p***

To estimate the relative expression of *LEP* and *miR-9-5p* in placentae, these primers were used for quantitative real-time PCR (qRT-PCR):

*LEP*: CACACGCAGTCAGTCTCCTC (forward), GGCCAGCACGTGAAGAAGA (reverse)

*miR-9-5p*: GGGTCTTTGGTTATCTAGC (forward), CGTATCCAGTGCTCATAC (reverse)
